# Mapping glymphatic solute transportation through the perivascular space of hippocampal arterioles with 14 Tesla MRI

**DOI:** 10.1101/2023.09.14.557634

**Authors:** Xiaoqing Alice Zhou, Weitao Man, Xiaochen Liu, Sangcheon Choi, Yuanyuan Jiang, David Hike, Lidia Gomez Cid, Changrun Lin, Maiken Nedergaard, Xin Yu

## Abstract

The perivascular space (PVS) plays a crucial role in facilitating the clearance of waste products and the exchange of cerebrospinal fluid and interstitial fluid in the central nervous system. While optical imaging methods identify the glymphatic transport of fluorescent tracers through PVS of surface-diving arteries, their limited depth penetration impedes the study of glymphatic dynamics in deep brain regions. In this study, we introduced a novel high-resolution dynamic contrast-enhanced MRI mapping approach based on single-vessel multi-gradient-echo methods. This technique allowed the differentiation of penetrating arterioles and venules from adjacent parenchymal tissue voxels and enabled the detection of Gd-enhanced signals coupled to PVS of penetrating arterioles in the deep cortex and hippocampus. By directly infusing Gd into the lateral ventricle, we eliminated delays in cerebrospinal fluid flow and focused on PVS Gd transport through PVS of hippocampal arterioles. The study revealed significant PVS-specific Gd signal enhancements, shedding light on glymphatic function in deep brain regions. These findings advance our understanding of brain-wide glymphatic dynamics and hold potential implications for neurological conditions characterized by impaired waste clearance, warranting further exploration of their clinical relevance and therapeutic applications.

## Introduction

The glymphatic system facilitates the dynamic interchange of cerebrospinal fluid (CSF) and interstitial fluid (ISF), contributing to maintaining regular physiological equilibrium and effectively removing interstitial substances^1–4^. Among the various compartments encompassing the intracranial interstitial extracellular space, the perivascular space (PVS) between the astrocyte endfeet and the parenchymal perforating vessels has been proposed to play a crucial role underlying the glymphatic clearance^5–9^. Two-photon optical imaging studies have well described the glymphatic cerebrospinal fluid circulation through the PVS from the surface diving arteries^10^. And, the latest work has shown the glymphatic flow through PVS of surface diving arteries can be further regulated by neuronal activity of the adjacent cortex^11^. However, given the limited penetrating depth of the optical imaging method, the dynamic changes of glymphatic flow in the deep cortex or subcortical regions remain to be elucidated.

Magnetic resonance imaging (MRI) has been applied to detect the PVS in human brains^12,13^, in particular, to detect the enlarged PVS as a potential marker of brain dementia^14^. To measure the potential glymphatic flow through PVS, the Gd-based dynamic contrast-enhanced (DCE) MRI has been applied to map both animal and human brains^12–24^. Through either intracisternal or intraventricular delivery of the Gd agent, the PVS-specific signal enhancement is also mainly detected from the large diving arteries, e.g. the middle cerebral artery (MCA) or through the posterior cerebral artery (PCA) supplying the hippocampus^25^. Most of the rodent DCE MRI studies have applied preclinical high-field MRI scanner at 9.4T or higher to ensure sufficient signal-to-noise ratio (SNR) up to 100um isotropic resolution. However, the measurement of solvent transportation through PVS of micro-vessels penetrating the deep cortex or subcortical brain regions with 20-50µm diameters remains challenging given the limited spatial resolution.

In this study, we developed a novel DCE MRI approach based on the single-vessel multi-gradient-echo (MGE) imaging scheme^26,27^ to measure Gd-enhanced PVS in mouse brains using a 14T preclinical MRI scanner. To ensure sufficient SNR with high spatial resolution images (50um isotropic resolution for the whole brain or 50×50×250um for temporal dynamic mapping at 10s TR), we implanted a radiofrequency (RF coil) above the skull with a micro-capillary targeting the lateral ventricle of the mouse brain for real-time DCE-MRI. This high-resolution DCE method enabled the differentiation of vessel voxels from peri-vessel voxels to verify the Gd-enhanced signals in PVS of penetrating arterioles of the cortex and hippocampus of mouse brains.

## Materials and Methods

### Animal surgical procedures

All animal surgical and experimental procedures were conducted following the Guide for the Care and Use of Laboratory Animals and approved by the Massachusetts General Hospital Subcommittee on Research Animal Care (Protocol No. 2020N000073). We used 10 healthy 6-month-old female C57/BL6 mice (The Jackson Laboratory, Maine, United States), 6 for perivascular space distribution, and 4 for glymphatic flow dynamics. Craniotomies were made above the right lateral ventricle (LV) under the microscope. 100nm/g Gd-filled capillary tube (Silica, Polyimide Coated Smooth Solid Tubing, 0.24mm, Digikey, MN), pre-connected to a micro-syringe (10 µL, WPI, FL) at one end, was inserted into the LV with the following coordinates: caudal 0.59mm, lateral 1.25mm, ventral 1.85mm from bregma. The mice then underwent craniotomy surgery to install an MRI coil, and a customed MRI coil was positioned over the cortex and tested for tunning and matching signal with a network analyzer. The coil ring was lifted ∼0.5mm above the surface of the skull and held in place while a thin layer of 404 cyanoacrylate glue was applied to connect the skull with the coil. Once dried (∼8 mins), 2part dental cement (Dental Cement Kit, Stoeling, IL) was mixed and applied to cover the coil and exposed bone paying special note to the base of the coil to firmly secure it and avoiding drips into the eye. The edges of the skin were then attached with 454 cyanoacrylate glue. After the dental cement had fully hardened (∼10 mins), the mouse was released from the stereotaxic stage and received subcutaneous injections of Dexamethasone and Cefazolin. During surgical sessions, mice were anesthetized at the same mixture of medical air (70 mL/min), oxygen (15 mL/min), and isoflurane (2%). A heating pad was used to ensure the mouse’s body temperature was maintained near 37° C throughout anesthesia. A PE10 tube was inserted into the lateral tail veins of the mice for Gd intravenous infusion.

### Magnetic Resonance Imaging

After the animal was secured in an MRI holder with ear and tooth bars, isoflurane was then reduced to 1.5%. The respiration rate and rectal temperature were continuously monitored (Model 1030, SAII, USA), with the temperature maintained at 36.5-37C by an air heater. The physiology was maintained within normal ranges, with a respiration rate of 80-110 breaths per minute throughout the MRI scans.

MRI was obtained by using a 14T/13 cm horizontal-bore magnet (Magnex Scientific) interfaced through the Bruker AV Neo system (Bruker). The scanner has a 6 cm gradient set with a strength of 100 Gauss per cm (G/cm) and a 150 μs rise time, as well as 6 channels up to 2^nd^/3^rd^ order shimming (Resonance Research Inc.). An implantable RF transceiver surface coil with an outer diameter of 9.5 mm (Mribot Inc.) was used for image acquisition.

To examine whether the perivascular space of deep cortical layers and subcortical regions is involved in the glymphatic flow circulation, we apply the ultra-high resolution MGE sequence (50um isotropic) to map differential Gd-enhanced signals between intravenous and intraventricular Gd administration. A 3D MGE sequence was applied with the following parameters: TR: 50 ms; TE: 2, 4.5, 7, 9.5 ms; flip angle: 85°; matrix: 256 × 256 x 160; in-plane resolution: 50 μm isotropic. Scan time: 1h 30min.We first acquired blood Gd-based flow maps with MGE MRI (tail-vein Gd infusion, 12nl/min for 0.1-0.2ml for 90 min), which highlights all vessel voxels as vascular control maps. Secondly, we acquired another set of MGE images with the intraventricular Gd infusion (20nl/min, for 1-2ul for 90 min).

Besides mapping the accumulative glymphatic transportation with whole brain MGE, we acquired DCE MGE images with ultra-high resolution to cover both the hippocampus of the mice. A 2D MGE sequence was applied with the following parameters: TR: 80 ms; TE: 2.5, 5.5, 8.5, 11.5ms; flip angle: 85°; matrix: 256 × 256; in-plane resolution: 50 × 50 μm^2^; slice thickness: 250 µm. Scan cycle:13.68s, repetition:120. Scan time: 27min22s. MGE images were acquired with the intraventricular Gd infusion (100nl/min, 1ul for 10 min) starting at 21st cycle.

## Results

We implemented the ultra-high resolution MGE sequence to characterize two aspects of glymphatic solvent transport through PVS.

First, the 3D whole brain Gd-enhanced MR images were acquired with 50µm isotropic resolution using the MGE sequences in two conditions: intravenous infusion of Gd (Gd_iv_) and intraventricular infusion of Gd (Gd_ivtr_) consecutively (**Fig 1A**). 3D images with Gd_iv_ detected some penetrating vessels both in the cortex and subcortical regions, e.g., hippocampus. After intraventricular infusion of Gd, there is a clear signal enhancement in the parenchyma regions near the ventricles. To verify the potential Gd-enhanced PVS, three penetrating vessels in the cortex were identified based on the Gd_iv_ condition. Fig 1B showed MGE images at the first and third TE, enabling the differentiation of arterioles from venules^26^. The deoxygenated blood in venules at longer TE, i.e., 3^rd^ TE, led to faster T2* decay to dimmish the venule signal despite the Gd effect. **Fig 1C** showed the line profiles around the three penetrating vessels, demonstrating the three vessels with high signal intensity at Gd_iv-TE1_ (**Fig 1B**, colored overlap image) but only two arterioles remained with high signal intensity at Gd_iv-TE3._ Meanwhile, we also plot the line profile of the Gd_ivtr_, presenting signal enhancement located at the two arterioles but with a slighter wider distribution (**Fig 1C**). Similarly, multiple penetrating vessels through the hippocampus of the two hemispheres were also identified in MGE images based on the Gd_iv_ condition (**Fig 1D**). Line profile plots of Gd_ivtr_ also showed a wider spatial distribution of signal enhancement near the hippocampal vessels (**Fig 1E, F**). **Fig 1G** showed the averaged vessel plots centered at the peak signal, showing a slightly wider distribution for the Gd_ivtr_, indicating the potential PVS-related Gd enhancement. It should be noted that no signal enhancement was detected near venules in the Gd_ivtr_ condition, indicating that glymphatic Gd transport is less affected by the blood Gd.

**Figure 1.**
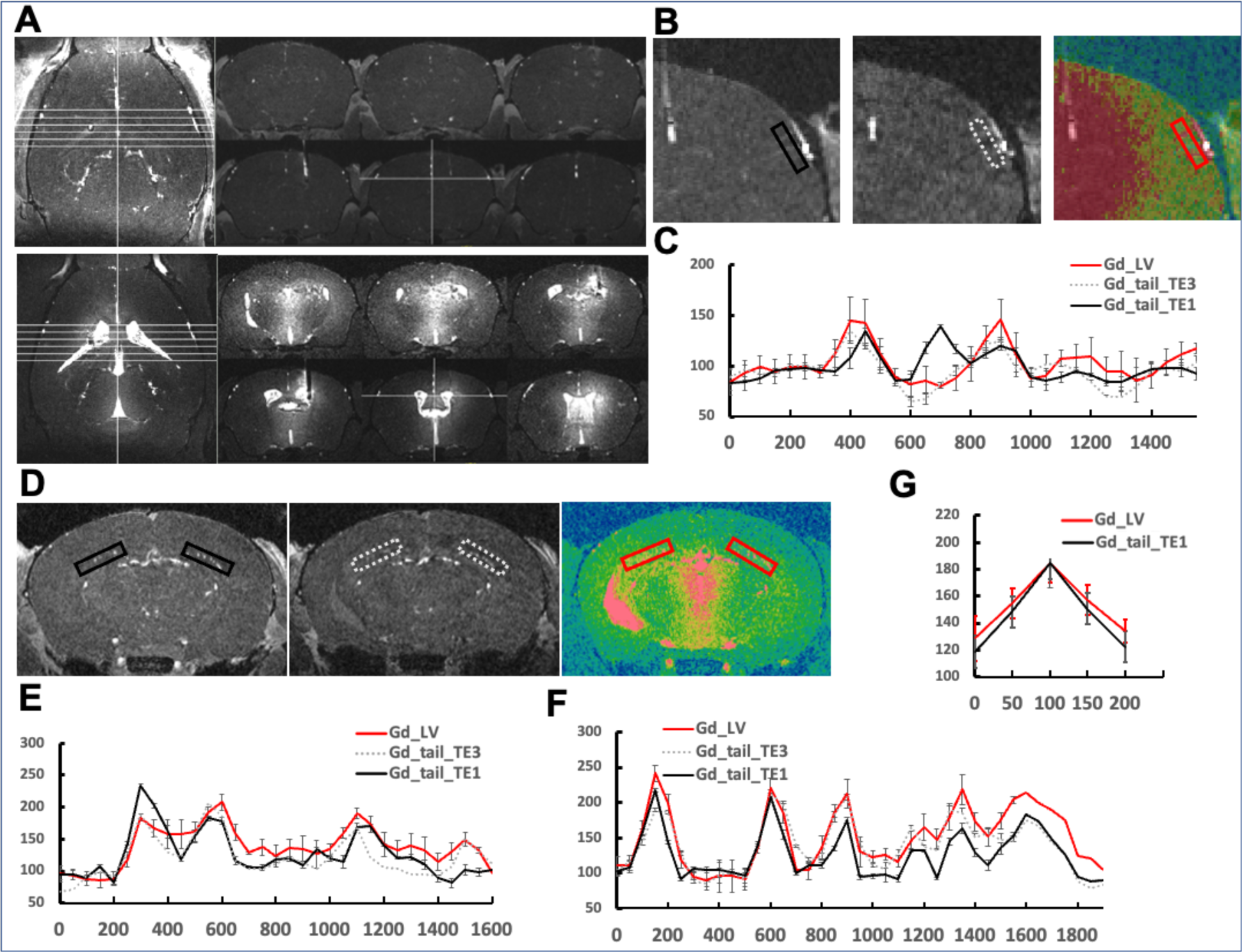
Detailed Analysis of Perivascular Space Distribution. **A,** Gd-enhanced single-vessel maps (upper panel) and Gd-enhanced CSF maps (lower panel), offering a comprehensive view of cerebral fluid dynamics within specific cortical regions. **B,** Representative slices of cortical vessels visualized with short (left), showing all vessels, and long TE (middle), which diminishes the visibility of venules due to faster T2* decay, but not arterioles, alongside a Gd-enhanced CSF map (right) that overlays the same area, facilitating a comparative analysis of solute distribution. **C,** The line profiles of vessels that show wider Gd distribution in Gd-enhanced CSF maps (red) compared to Gd-enhanced single-vessel maps, indicating a significant solute exchange within the perivascular space. **D,** Single-vessel maps of hippocampal vessels with varying echo times, emphasizing the differential visibility of vascular structures under different imaging conditions. **E&F,** The comparative line profiles of vessels (E, left), which indicate a broader Gd distribution space in Gd-enhanced CSF maps (red) compared with Gd-enhanced single-vessel maps, providing evidence of active solute transport through the perivascular spaces. **G**, The line profiles of the average of 10 vessels in the regions of interest in Panels B and D, synthesizing the spatial dynamics of Gd distribution and its implications for understanding cerebral solute transport mechanisms.

With this methodology, Gd-enhanced PVS mapping was conducted alongside the inflow vascular map of mouse brains. Extensive Gd signal enhancement suggests effective contrast penetration and distribution within the PVS, especially around major cerebral arteries as shown in **Fig 2A. Figures 2B and 2C** provide focused insights into the midbrain and pontine areas, respectively, with insets detailing the Gd enhancement patterns. Similarly, Figures 3 and 4 display the PVS in the cortex and hippocampus, respectively, highlighting the efficacy of MRI in capturing the intricate PVS architecture.

**Figure 2.**
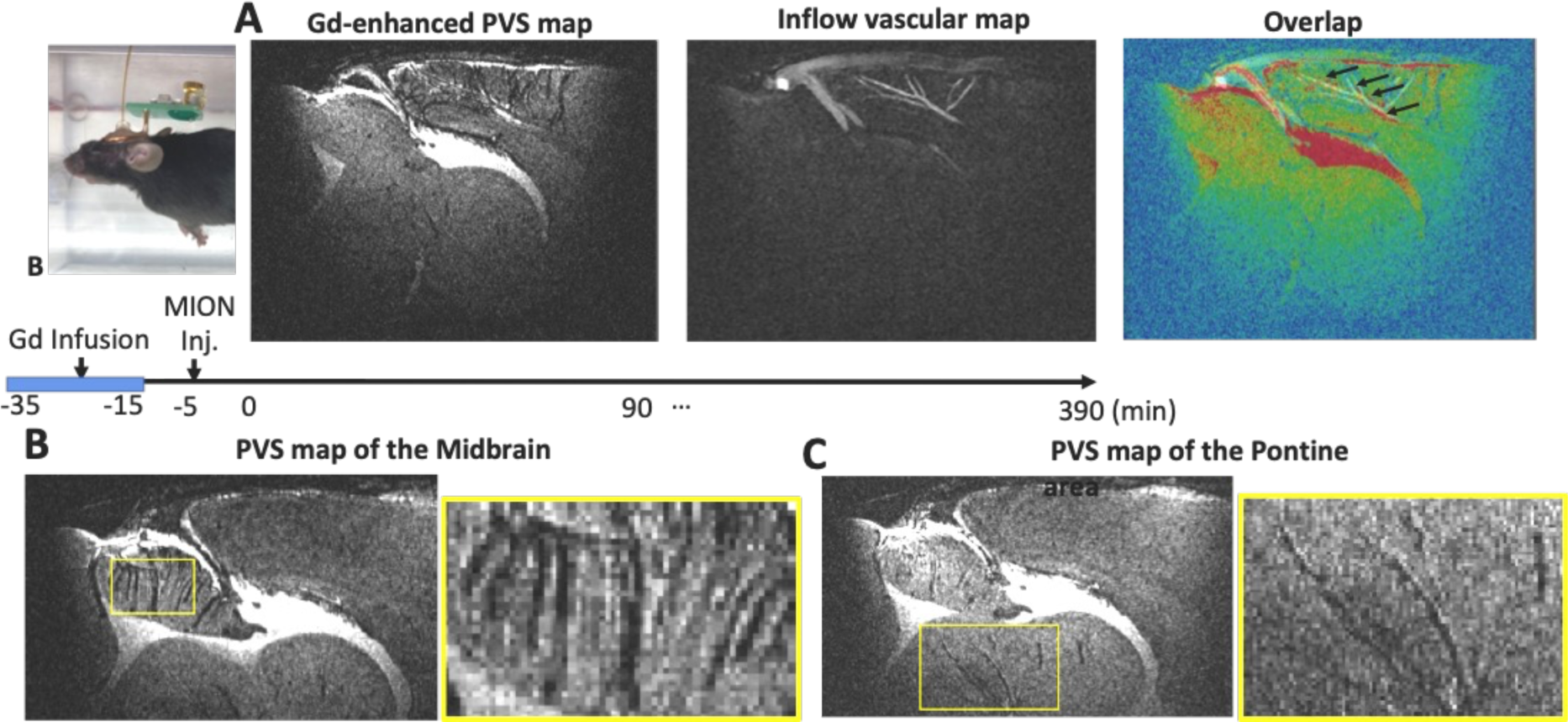
Visualization of PVS across major branches of anterior cerebral artery and other artery branches in the midbrain and pontine. A, Registration of the vascular map with a Gd-enhanced PVS map, displaying detailed imaging of PVS in major branches of the ACA. The colored overlays in Panel A depict the alignment of PVS with vascular structures, underscoring the effectiveness of Gd in enhancing these vital neuroanatomical features for detailed study. B&C, Representative slices that highlight the intricate structure of the PVS within the midbrain (B) and pontine areas (C).

**Figure 3.**
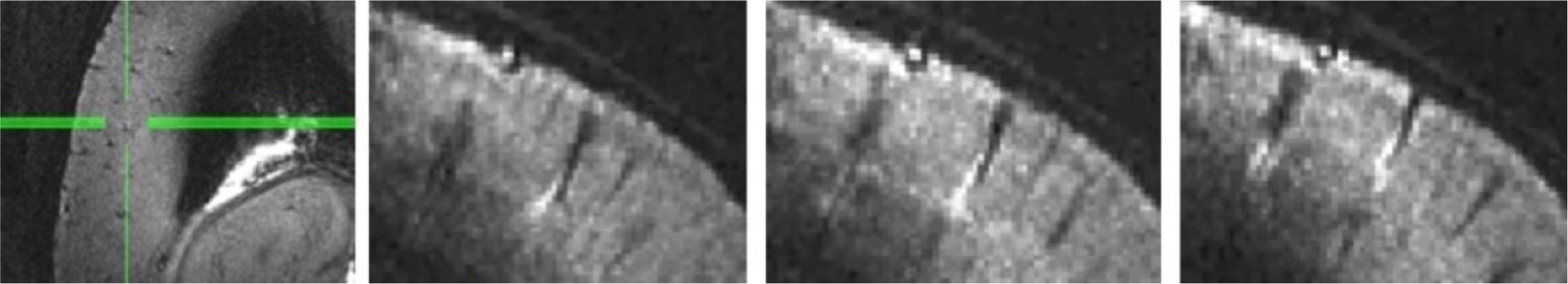
Visualization of PVS of penetrating arteries in the cortex. Representative slice of PVS in the cortical penetrating arterials, axial view and coronal view. Each image captures a distinct segment along the length of these vessels.

**Figure 4.**
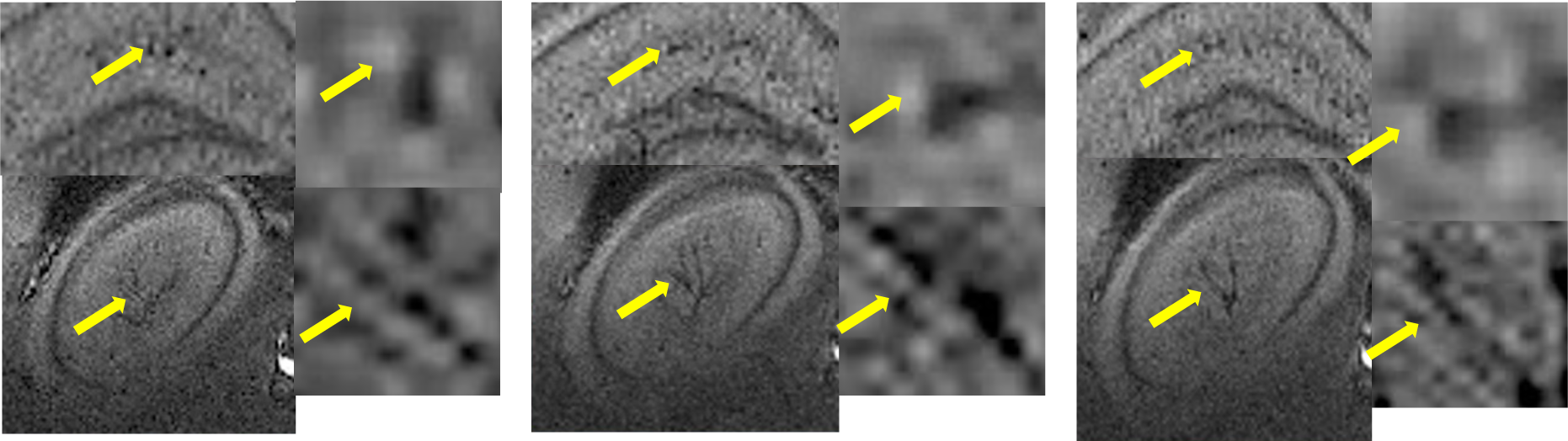
Visualization of PVS in penetrating hippocampal vessels. Representative MRI slices focusing on the PVS associated with the penetrating hippocampal vessels. Each image captures a distinct segment along the length of these vessels, demonstrating the glymphatic transport pathways as highlighted by the yellow arrows.

Moreover, we acquired higher-resolution 2D Gd-enhanced MR images at 20×20µm (**Fig 5A&B**) and 15×15µm (**Fig 5C**) in-plane resolutions. In the inflow maps, vessels were highlighted by bright signals, which were subsequently dampened post-iron injection and re-enhanced following Gd infusion, providing a clearer visualization of the PVS.

**Figure 5.**
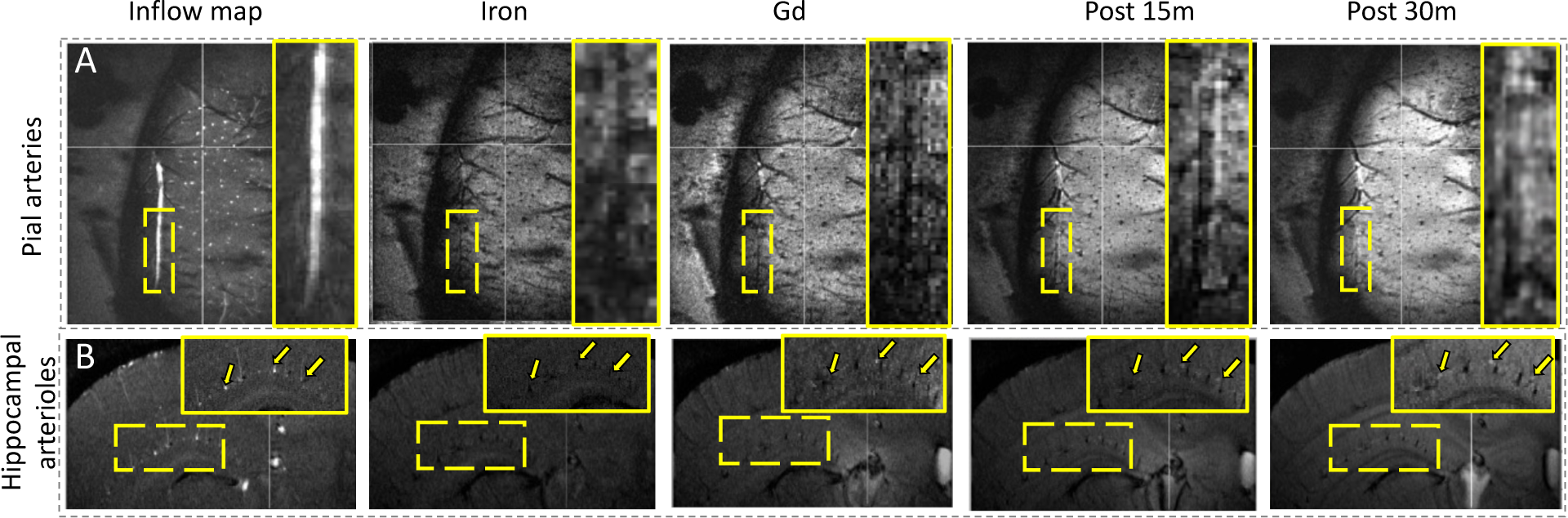
Perivascular space distribution detected by super resolution MRI. ***A,** PVS of pial arteries* branches with a resolution of 20×20×250 µm, where highlighted areas indicate the location of vessels. ***B**, PVS of penetrating hippocampal arterioles* with a resolution of 20×20×250 µm, where highlighted areas and arrows indicate the location of vessels. In both panels, highlighted areas and arrows point to specific vascular structures. From the inflow maps, vessels are initially marked by bright signals. After an intravenous iron injection, the vessel signals were dampened, but following Gd infusion, there is a marked enhancement within the PVS, evident from the enhanced signal visibility.

Further, slice-selective 2D MGE images with 50×50 µm in-plane resolution were obtained, oriented perpendicularly to the hippocampal penetrating vessels ^28^ (**Fig 6A**). High-resolution imaging facilitated the differentiation of vessel, peri-vessel, and adjacent parenchymal tissue voxels (**Fig 6B**) The Gd-enhanced DCE signals from three different ROIs plotted over time, before and after a 10-minute Gd infusion, showed a notable early signal increase in the peri-vessel ROIs compared to the adjacent parenchymal tissue. A slight signal drop in the vessel ROI could be attributed to T2* shortening caused by blood Gd during the infusion. This result conclusively demonstrates the PVS-specific glymphatic Gd transport in hippocampal penetrating vessels.

**Figure 6.**
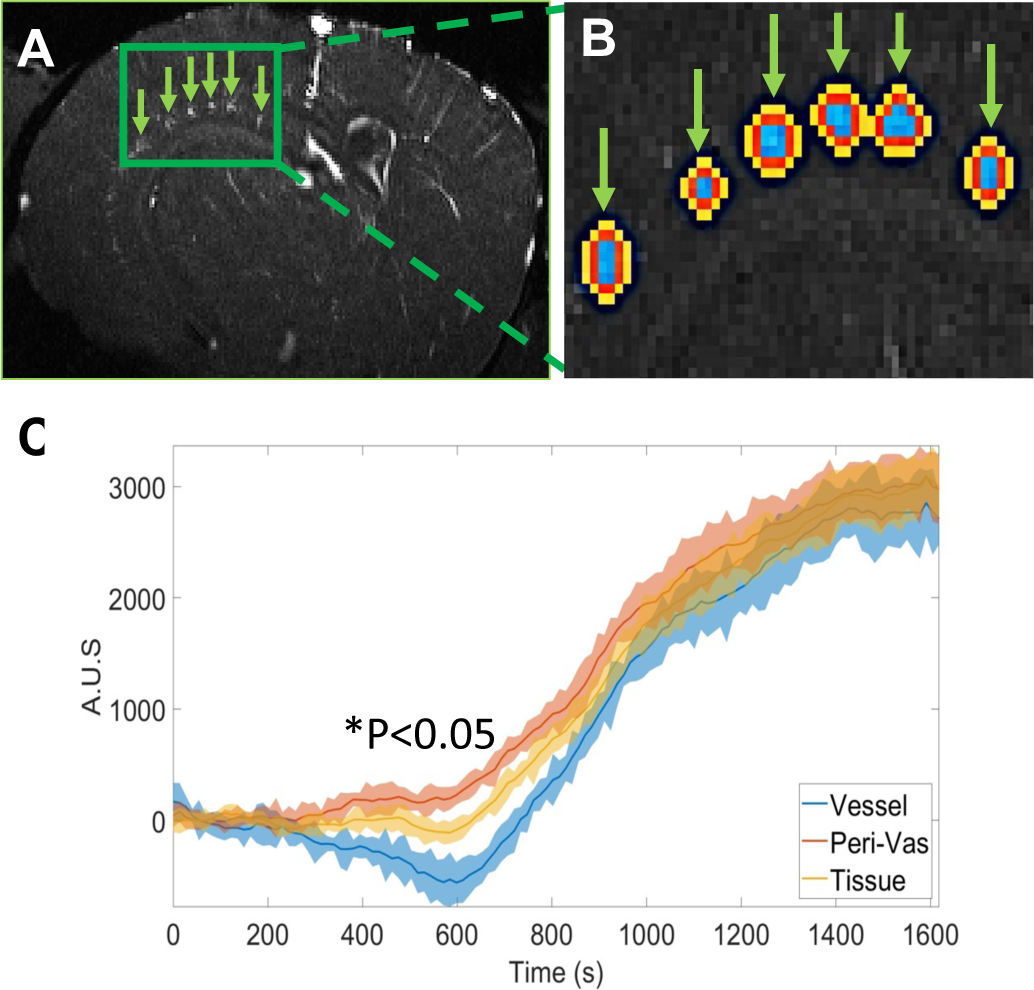
Dynamic Visualization of Hippocampal Perivascular Influx. **A**, A vessel-specific hippocampal map, where arrows pinpoint the identified vessels within the hippocampus, highlighted for clarity. **B**, ROIs colored to distinguish between vessels (blue), perivascular space (red), and surrounding tissue (yellow), aiding in the visualization of spatial relationships and interactions. **C**, A graphical representation of the time courses of Gd signal change across these ROIs, reflecting the differential dynamics in vessels, perivascular space, and surrounding tissue. The data points, accumulated from two trials per mouse across four mice, demonstrate significant statistical differences (*P<0.05), indicating distinct signal enhancements and temporal patterns in each specified region.

## Discussion

The present study aims to investigate whether glymphatic transport occurs in PVS of penetrating vessels in the deep brain regions. In contrast to the optical imaging methods that provide high spatial resolution to identify microscopic measurements of PVS from diving arteries primarily located at the cortical surface, Gd-based DCE MRI has been used to measure the brain-wide glymphatic flow dynamic changes. There are two ongoing challenges related to this issue with MRI. First is the limited spatial resolution to distinguish vessels from PVS and adjacent parenchymal tissues with conventional MRI methodology. Second is the difficulty in differentiating the diffusion through extracellular matrix versus glymphatic PVS-specific solvent transportation.

We tackled this problem by implementing a novel DCE MRI mapping scheme based on single-vessel MGE methods, originally developed to map hemodynamic responses from penetrating arterioles and venules. The MGE sequence enables image acquisition at different echo times (TE). By setting a large flip angle with a short time of repetition (TR), we are able to highlight certain penetrating vessels based on their orientation to the MRI slices. Also, due to the different T2* values of the oxygenated and deoxygenated blood in arterioles and venules, it also offers a possibility to distinguish arterioles from venules. When Gd is infused into blood, it can further enhance the vessel detection at shorter TE, and leads to faster T2* decay of venules. Thus, the MGE method offered an ideal image scheme to detect the penetrating micro-vessels with unique orientation with regard to the geometrical alignment of the MRI image field of view. It should be also noted that the hippocampal penetrating vessels align parallel to the cortical surface. This alignment is hard to detect by functional ultrasound based on the Doppler effect with a probe directly attached to the mouse head. Whereas, as a less sensitive method in comparison to the optical or acoustic imaging methods, it is critical to ensure sufficient SNR when mapping the micro-penetrating vessels with high spatial resolution and temporal resolution. We also implanted an RF coil to be attached to the skull, which enables a significant signal increase^29^. Given Gd-driven T1 shortening, it is practical to achieve ultra-high spatial resolution images with 30-50um to measure PVS-related signal changes near arteries and venules in deep brain regions. It should also be noted that the actual PVS near arteries with a diameter of 20-30µm is around 1-2µm^30^. The Gd-enhanced signal in peri-vessel voxels is still mainly due to partial volume effects from PVS. Importantly, Fig 2 showed the opposite sign of signal changes in vessel voxels and peri-vessel voxels indicating the PVS-specific contribution to signal increase in the peri-vessel voxels despite the relatively large voxel size.

In contrast to most of the Gd-based DCE MRI with intracisternal administration, we directly infused Gd into the lateral ventricle and focused on measuring PVS Gd transport in the hippocampal vessels. This administration scheme also avoided the time delay of CSF flow from the cisterna magna to the other ventricles, which can confound with the direct interstitial diffusion through the extracellular matrix. It has been shown that the low-frequency oscillation of arteriole diameter changes (i.e., vasomotion) can alter the glymphatic flow rate, leading to more efficient solvent transport through PVS than molecule dispersion-based diffusion^15,31^. The increased transportation rates are brain state dependent and vary from single-digit to double-digit percentage increase from quiet wakefulness to sleep. The little difference further complicates the measurement of PVS-specific Gd signal enhancement of penetrating vessels from adjacent parenchymal tissues.

The 2D MGE slice was positioned to measure the cross-section of the hippocampal penetrating arteries that were around 1mm away from the lateral ventricles. Previous optical imaging studies have estimated a 10-30um/s glymphatic flow rate through peri-arteriole PVS based on 70-2000kDa solutes. And given the obstructive scale in the PVS, the actual transportation rate was further slowed down to be around 40-70s per 100µm distance. In our study, the Gd-DTPA is near 1kDa, which is much smaller than the fluorescent tracers used for optical imaging. The PVS-specific glymphatic transportation showed significant signal enhancement around 140s, enabling the estimate of 7s per 100 µm distance. We will further verify the Gd-DTPA-specific diffusion through the interstitial fluid of the extracellular matrix, which can be compared with the PVS-specific Gd transportation. It should also be noted that the estimated glymphatic flow rate in venule PVS is much slower, which may not be sufficient to be differentiated from the parenchymal diffusion effect.

The results from our methodology demonstrate not only the practical utility of Gd-enhanced PVS mapping in revealing the dynamics of glymphatic transport but also emphasize the robustness of our imaging technique in differentiating the vascular and perivascular architectures within the mouse brain. The distinct Gd signal enhancements observed around the major cerebral arteries and within specific brain regions such as the midbrain, pontine area, cortex, and hippocampus, underscore the precision of this imaging modality in mapping the perivascular pathways that are crucial for validating the brain-wide existence of glymphatic transporation. Furthermore, the use of higher-resolution 2D Gd-enhanced MR images has significantly refined our ability to visualize and analyze the PVS architecture through the hippocampal structure, enabling deep-brain region-specific characterization of altered solute transportation through the PVS during brain degeneration in future studies. To be noted, it is evident that the visualization of PVS surrounding the micro-vessels that extend across various orientations with relevance to the magnetic z_0_ directions. The dipole effect in MRI due to the iron-induced susceptibility changes is mostly predominant in the z_0_-direction of the magnet, presenting altered phase signals due to the field inhomogeneity changes along the z_0_ axis. In contrast, the PVS-induced signal gain near mivro-vessels captured in the high-resolution MRI images are aligned in multiple spatial directions, which is not simply caused by the dipole effect. To further verify this potential confounding issues, we will analyze the phase maps from the high-resolution MRI images with Gd-enhanced signals from PVS of mice aligned with different orientation in relevance to the z_0_ direction.

In conclusion, this study represents a step forward in the field of glymphatic research by mapping the perivascular pathway and dynamics through micro-vessels penetrating deep brain regions, e.g., the hippocampus, of the mouse brain. The findings contribute to our understanding of glymphatic function and have the potential to shed light on neurological conditions where impaired waste clearance is implicated. Further research is warranted to explore the clinical implications and therapeutic potential of these discoveries.

## Acknowledgement

This research was funded by Alzheimer’s association (AARFD-23-1145375), NIH funding (RF1NS113278, RF1NS124778, R01NS122904, R21NS121642,), NSF grant 2123971, and the S10 instrument grant (S10 MH124733–01) to Martinos Center.

## Notes

### Competing Interest Statement

The authors have declared no competing interest.

### Summary of Updates

This version of the manuscript has been revised to update the following in the Results section: 1. Ultra-High Resolution MGE Sequence Implementation: Detailed the application of the ultra-high resolution multi-gradient-echo (MGE) sequence to characterize glymphatic solute transport through perivascular space (PVS). 2. Hippocampal Penetrating Vessels: Detailed the identification of multiple penetrating vessels in the hippocampus using MGE imaging. 3. Higher-Resolution 2D Gd-Enhanced MR Imaging: Introduced higher-resolution 2D Gd-enhanced MR images at 20×20um and 15×15um in-plane resolutions. Overall, these revisions enhance the clarity and depth of the Results section, providing a comprehensive and detailed analysis of the glymphatic solute transport dynamics through the PVS in the hippocampus and other brain regions. The updated data and visual representations underscore the robustness and precision of the high-resolution MRI techniques used in this study.

## References

1 Benveniste, H. et al. The Glymphatic System and Waste Clearance with Brain Aging: A Review. Gerontology 65, 106–119 (2019). 10.1159/000490349

2 Da Mesquita, S. et al. Functional aspects of meningeal lymphatics in ageing and Alzheimer’s disease. Nature 560, 185–191 (2018). 10.1038/s41586-018-0368-8

3 Kress, B. T. et al. Impairment of paravascular clearance pathways in the aging brain. Annals of neurology 76, 845–861 (2014). 10.1002/ana.24271

4 Zeppenfeld, D. M. et al. Association of Perivascular Localization of Aquaporin-4 With Cognition and Alzheimer Disease in Aging Brains. JAMA Neurol 74, 91–99 (2017). 10.1001/jamaneurol.2016.4370

5 Marina, N. et al. Astrocytes monitor cerebral perfusion and control systemic circulation to maintain brain blood flow. Nature communications 11, 131 (2020). 10.1038/s41467-01913956-y

6 Kisler, K., Nelson, A. R., Montagne, A. & Zlokovic, B. V. Cerebral blood flow regulation and neurovascular dysfunction in Alzheimer disease. Nature Reviews Neuroscience 18, 419–434 (2017). 10.1038/nrn.2017.48

7 MacVicar, B. A. & Newman, E. A. Astrocyte regulation of blood flow in the brain. Cold Spring Harb Perspect Biol 7 (2015). 10.1101/cshperspect.a020388

8 Iliff, J. J. et al. A paravascular pathway facilitates CSF flow through the brain parenchyma and the clearance of interstitial solutes, including amyloid β. Sci Transl Med 4, 147ra111 (2012). 10.1126/scitranslmed.3003748

9 Attwell, D. et al. Glial and neuronal control of brain blood flow. Nature 468, 232–243 (2010). 10.1038/nature09613

10 Iliff, J. J. et al. A paravascular pathway facilitates CSF flow through the brain parenchyma and the clearance of interstitial solutes, including amyloid beta. Sci Transl Med 4, 147r–a111 (2012). 10.1126/scitranslmed.3003748</otherinfo>

11 Holstein-Rønsbo, S. et al. Glymphatic influx and clearance are accelerated by neurovascular coupling. Nature neuroscience 26, 1042–1053 (2023). 10.1038/s41593-023-01327-2

12 Ringstad, G. et al. Brain-wide glymphatic enhancement and clearance in humans assessed with MRI. JCI Insight 3 (2018). 10.1172/jci.insight.121537

13 Eide, P. K. & Ringstad, G. MRI with intrathecal MRI gadolinium contrast medium administration: a possible method to assess glymphatic function in human brain. Acta Radiol Open 4, 2058460115609635 (2015). 10.1177/2058460115609635

14 Harrison, I. F. et al. Impaired glymphatic function and clearance of tau in an Alzheimer’s disease model. Brain : a journal of neurology 143, 2576–2593 (2020). 10.1093/brain/awaa179

15 Iliff, J. J. et al. Cerebral arterial pulsation drives paravascular CSF-interstitial fluid exchange in the murine brain. The Journal of neuroscience : the official journal of the Society for Neuroscience 33, 18190–18199 (2013). 10.1523/jneurosci.1592-13.2013

16 Gaberel, T. et al. Impaired glymphatic perfusion after strokes revealed by contrast-enhanced MRI: a new target for fibrinolysis? Stroke; a journal of cerebral circulation 45, 3092–3096 (2014). 10.1161/strokeaha.114.006617

17 Gakuba, C. et al. General Anesthesia Inhibits the Activity of the “Glymphatic System”. Theranostics 8, 710–722 (2018). 10.7150/thno.19154

18 Mestre, H. et al. Aquaporin-4-dependent glymphatic solute transport in the rodent brain. Elife 7 (2018). 10.7554/eLife.40070

19 Koundal, S. et al. Optimal Mass Transport with Lagrangian Workflow Reveals Advective and Diffusion Driven Solute Transport in the Glymphatic System. Scientific reports 10, 1990 (2020). 10.1038/s41598-020-59045-9

20 Mortensen, K. N. et al. Impaired Glymphatic Transport in Spontaneously Hypertensive Rats. The Journal of neuroscience : the official journal of the Society for Neuroscience 39, 6365–6377 (2019). 10.1523/jneurosci.1974-18.2019

21 Hadjihambi, A. et al. Impaired brain glymphatic flow in experimental hepatic encephalopathy. Journal of hepatology 70, 40–49 (2019). 10.1016/j.jhep.2018.08.021

22 Ding, G. et al. MRI investigation of glymphatic responses to Gd-DTPA infusion rates. J Neurosci Res 96, 1876–1886 (2018). 10.1002/jnr.24325

23 Ohno, K. et al. Kinetics and MR-Based Monitoring of AAV9 Vector Delivery into Cerebrospinal Fluid of Nonhuman Primates. Mol Ther Methods Clin Dev 13, 47–54 (2019). 10.1016/j.omtm.2018.12.001

24 Lee, H. et al. Quantitative Gd-DOTA uptake from cerebrospinal fluid into rat brain using 3D VFA-SPGR at 9.4T. Magnetic resonance in medicine : official journal of the Society of Magnetic Resonance in Medicine / Society of Magnetic Resonance in Medicine 79, 1568–1578 (2018). 10.1002/mrm.26779

25 Koundal, S. et al. Optimal Mass Transport with Lagrangian Workflow Reveals Advective and Diffusion Driven Solute Transport in the Glymphatic System. Scientific reports 10, 1990 (2020). 10.1038/s41598-020-59045-9

26 Yu, X. et al. Sensory and optogenetically driven single-vessel fMRI. Nature methods 13, 337–340 (2016). 10.1038/nmeth.3765

27 He, Y. et al. Ultra-Slow Single-Vessel BOLD and CBV-Based fMRI Spatiotemporal Dynamics and Their Correlation with Neuronal Intracellular Calcium Signals. Neuron 97, 925–939 e925 (2018). 10.1016/j.neuron.2018.01.025

28 Chen, X. et al. Mapping optogenetically-driven single-vessel fMRI with concurrent neuronal calcium recordings in the rat hippocampus. Nature communications 10, 5239 (2019). 10.1038/s41467-019-12850-x

29 Chen, Y. et al. Focal fMRI signal enhancement with implantable inductively coupled detectors. NeuroImage 247, 118793 (2022). 10.1016/j.neuroimage.2021.118793

30 Bojarskaite, L. et al. Sleep cycle-dependent vascular dynamics in male mice and the predicted effects on perivascular cerebrospinal fluid flow and solute transport. Nature communications 14, 953 (2023). 10.1038/s41467-023-36643-5

31 van Veluw, S. J. et al. Vasomotion as a Driving Force for Paravascular Clearance in the Awake Mouse Brain. Neuron 105, 549–561.e545 (2020). 10.1016/j.neuron.2019.10.033

